# diempy: fast and reference-free genome polarisation

**DOI:** 10.64898/2026.02.18.706591

**Authors:** Derek Setter, Konrad Lohse, Stuart J.E. Baird

## Abstract

Most ancestry-assignment methods rely on putatively pure reference panels, which are often unrealistic and bias inference. The genome polarisation algorithm *diem*, introduced previously, avoids reference panels by jointly inferring the polarity of common allelic states and quantifying variant diagnosticity via an expectation–maximisation procedure. Here we present `diempy`, an efficient python implementation of *diem* coupled with tools that turn polarised calls into analysis-ready outputs. `diempy` offers lossless VCF-to-*diem* BED conversion; ploidy-aware handling of individuals and chromosomes; flexible masking of sites, regions and individuals; and interactive visualisation of polarised genomes, hybrid indices, clines and ternary plots. Post-processing functions include DI thresholding, kernel smoothing, and automatic detection and run-length encoding of contiguous ancestry tracts. BED-based I/O facilitates integration with population-genomic workflows (e.g. filtering by annotation or ploidy). These features make reference-free genome polarisation with `diempy` practical and reproducible for studies of population structure, admixture and species barriers.

## Introduction

Analyses aimed at understanding population structure, hybridisation and admixture often start by deconstructing and visualising the ancestry contributions in a sample of genomes. In the simplest case of two ancestry sources, individual diploid genomes are mosaics of tracts that are homozygous for either source or heterozygous by source. Previous ancestry assignment and chromosome paintings have relied on “ancestry informative variants” that are defined *a priori* using a set of reference samples assumed to be pure. The assumption of pure reference panels is both biologically implausible and biases inference [9]. Genome polarisation as implemented in an expectation maximisation (EM) algorithm *diem* co-estimates both the diagnosticity – quantified as a diagnostic index (DI) – and polarity of variants and avoids the need for reference panels. The core EM algorithm for genome polarisation is efficient, deterministic and has been implemented in `Mathematica`, python, and R, [3], with a current R CRAN package [1]. Genome polarisation has been used in analyses of whole genome sequence (WGS) and reduced representation data to investigate species barriers and adaptive variation that penetrates them across a wide range of systems [3, 8, 5, 21]. We stress that polarization in *diem* refers to the assignment of alleles with respect to the sides of a barrier and should not be confused with determining the ancestral state, for which other methods are available [18].

The flexibility and generality of reference-free genome polarisation avoids the need for any arbitrary pre-filtering of data and instead gives users the ability to explore and visualise polarised data for a range of diagnosticity thresholds and smoothing decisions.

Here we present `diempy`, an efficient python implementation of the *diem* genome polarisation algorithm. `diempy` includes a preprocessing module as well as functions for data visualisation, diagnosticity thresholding and smoothing – core post-processing steps – for which no tools were available previously.

Crucially, `diempy` uses bed files to specify input and output. This allows incorporation of genome polarisation into general population genomic analysis workflows that may involve excluding or masking data based on ploidy, functional annotation, quality filters or specific patterns of sequence variation.

## Methods

`diempy` implements the EM algorithm for genome polarisation of Baird et al 2023 [3] as an open source software package in Python 3 [23] (version ≥ 3.9). The source code is available on GitHub (https://github.com/Studenecivb/diemPy), and the package is available on PyPi for direct installation using `pip` (https://pypi.org/project/diempy/). Below, we outline the core functionality of `diempy`, and the three classes that implement the algorithm and subsequent analyses.

### Core Functionality

The package provides several main analysis workflows:

1. **Data Import and Processing**
  - Convert VCF files to *diem* format using `vcf2diem`
  - Read diem BED files with `read_diem_bed`
  - Handle ploidy information and masking of individuals and sites
2. **Polarization Analysis**
  - Initialize polarization using random null
  - Run EM algorithm to optimize variant polarization
  - Calculate diagnostic indices and support values
3. **Genome exploration**
  - Dynamic plots showing the influence of diagnostic index thresholding
  - Genome summaries at the level of individuals and chromosomes
  - Cline, ternary and iris plots
4. **Post-Polarization Analysis**
  - Compute hybrid indices for individuals
  - Apply thresholding to filter less informative variants
  - Perform kernel smoothing
  - Generate tracts of contiguous ancestry and store them as contigs
  - convert reference genome + BED intervals of `diempy` output to FASTA

### Core Classes and their Functions

There are three main classes that serve both as an efficient internal datastructure and provide easy access to the data for post-polarization analysis, namely the `DiemType`, `Contig` and `Interval` classes. The `DiemType` class contains the genotype data formatted for *diem* and results of the polarization. The `DiemType` stores a matrix of `Contigs`, one for each individual and chromosome in the dataset. Every `Contig` is composed of an ordered list of `Intervals`, each representing a contiguous tract of polarised ancestry which is run length encoded, i.e. defined by its start (the position of the first variant) and end (last variant). Below, we outline the main components and key methods of each class.

### DiemType Class

The `DiemType` is the central data structure that stores the polarised data for each chromosome. The Diem Matrix By Chromosome, **DMBC** is, a lossless encoding of genotype information. It contains individuals as rows and variants as columns: entries take token values in *{*0,1,2,3*}* corresponding to the *diem* genotypic states *{*U,0,1,2*}*: missing data, homozygous for the commonest allele, heterozygous, or homozygous for the second most common allele, respectively. Generalised over arbitrary ploidy the relevant terms become homogeneous vs heterogeneous for the two commonest alleles [3]. Other attributes of the `DiemType` are genomic positions in base pairs **posByChr**, chromosome lengths **chrLengths**, individual meta data including per-chromosome ploidy **ploidyByChr**. Most importantly, the `DiemType` stores polarization results including the initial null polarity, the final polarity, diagnostic indices (DI), and support values. The key methods for the `DiemType` include:

- `polarize()`: performs the expectation-maximization that polarises the data
- `apply_threshold()`: filters variants based on diagnostic index
- `smooth()`: applies the Laplacian kernel smoothing
- `sort()`: sorts the individuals (and corresponding data) by hybrid index
- `create_contig_matrix()`: builds the contigs from the polarised and processed data

### Contig Class

The `Contig` class represents a collection of genomic intervals (**intervals**) for a specific individual (**indName**) and chromosome (**chrName**), essentially describing the complete ancestry structure along that chromosome. The contig class contains two key methods. `printIntervals()` displays interval information in a human-readable format, and `get_intervals_of_state(state)` which returns all intervals with the specified state.

### Interval Class

The `Interval` class represents contiguous tracts of the genome with the same ancestry state for a given individual and chromosome. The attributes include the chromosome name (**chrName**), individual name (**indName**), the indices of the left- and right-most variants defining the interval, (**idxl** and **idxr**, inclusive) the corresponding genomic positions (**l** and **r**), as well as the state (**state**).

The key methods are `span()` which returns the physical span of the interval in base pairs, and `mapSpan(chrLength)` that returns the span relative to the specified total length of the chromosome (i.e. on a unit interval).

### Dependencies

`diempy` has the following dependencies: `numpy` ≥ 1.20.0 [10], `pandas` ≥ 1.3.0 [19, 22], `numba` ≥ 0.56.0 and ≤ 2.3 [17], `matplotlib` ≥ 3.4.0 [12], `docopt` ≥ 0.6.2 [14], `scikit-allel` ≥ 1.3.0 [20], `joblib` ≥ 1.5.3 [13], `ipympl` ≥ 0.10.0 [12], and `jupyterlab` ≥ 4.5.3 [16].

## Results

### Benchmarking

To benchmark the performance of `diempy`, we simulated a recent pulse of admixture between two divergent populations using `msprime` [15, 4] (see supplementary material). Note that while this demography is apropos of barrier analysis, the benchmarking results are model invariant, i.e. only the size of the dataset matters. We compared the run time for sequential (*k* = 1 core) and parallel processing on either *k* = 5 or *k* = 20 cores. We recorded the run time for a single iteration of the expectation maximization step of the *diem* algorithm for *n* = 10 (blue), *n* = 100 (magenta), and *n* = 1000 samples (red) as a function of the number of variants in the dataset (Fig. 1). Here, the variants are distributed evenly across 100 chromosomes.

**Figure 1:**
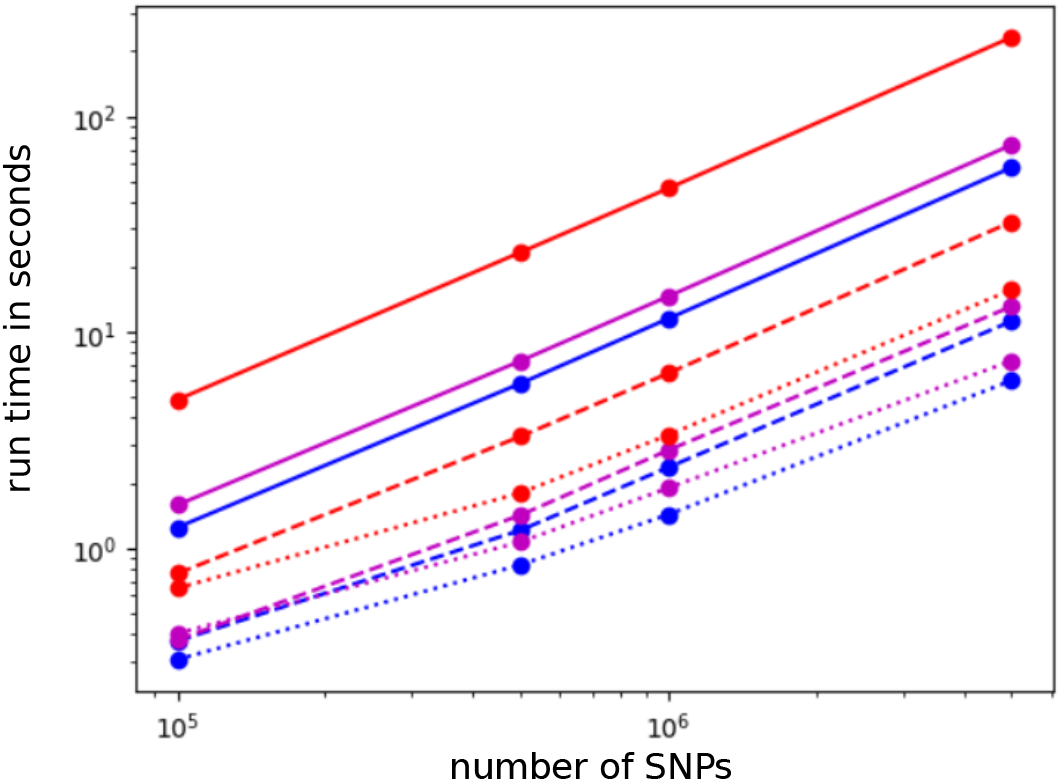
Run time for one iteration of the *diem* EM algorithm in seconds as a function of the number of variants for a sample of 10 individuals (blue), 100 individuals (magenta) and 1000 individuals (red). Solid lines represent sequential processing on a single core. The run time with parallel processing is shown for 5 and 20 cores (dashed and dotted lines respectively).

Run time increases linearly with the number of variants regardless of the number of samples or cores used (Fig. 1). However, we observe some overhead cost to parallel processing when the number of variants is small. In contrast, there is a sub-linear cost to increasing the number of samples: a 10-fold increase in the number of samples from *n* = 10 to *n* = 100 increases run time by approximately 1.2 fold, while a 100-fold increase in sample size (to *n* = 1000) induces only a 2.8 fold increase in run time.

Comparing parallel to sequential processing, we see that for realistically large datasets, *k* = 5 cores yield a 5-fold reduction in run time for *n* = 100 samples, as expected. Intriguingly, this improves to a 7-fold reduction for *n* = 1000 samples, possibly due to minor differences in the way memory is handled between the sequential and parallel versions of the algorithm. However, there are diminishing returns for parallel processing. Using *k* = 20 cores only provides a 10-fold decrease in run time for *n* = 100 and a 14-fold reduction for *n* = 1000 samples.

We also use the benchmarking data to test the memory requirements for `diempy`. For both the sequential and parallel versions of the algorithm, memory scales with the product of the number of variants and number of samples. For example, polarisation of *m* = 1, 000, 000 variants and *n* = 1, 000 individuals, i.e. 10^9^ data points, requires approximately 12GB of memory.

### Example Analysis

We provide an overview and extensive API documentation of the software (https://diempy.readthedocs.io), as well as an illustrative tutorial and workflow using an example dataset and jupyter lab (https://github.com/DerekSetter/tutorial-DiemPy/). We strongly recommend using the tutorial as a starting point for diempy analysis.

Here, we illustrate the step-by-step use of `diempy` using the example dataset provided with the tutorial. The example dataset comprises whole genome sequence (WGS) data for 20 individuals from a sister species pair of Scarce Swallowtail butterflies: *Iphiclides podalirius* and *I. feisthamelii*. The data includes individuals sampled throughout the range of each species as well as six individuals from the hybrid zone of the two species. Analyses of the full dataset (including *diem* polarisation) are described in detail in [5]. The example data comprise a subset of 2,000 consecutive variants from each of the 30 chromosomes (about 0.13% of the total data).

### Data and pre-processing

To perform genome polarization using `diempy`, the user first provides a vcf file that contains genotype data for all individuals and chromosomes, as well as the physical length of each chromosome in base pairs (in the header). The vcf input need not be aligned to a chromosome level reference: that is, we use ‘chromosome’ to mean known linkage group, and so as interchangeable with ‘scaffold’ or indeed ‘ddRAD interval’. It makes no difference to *diem* how variants are partitioned. The data are pre-processed for input into `diempy` using the **vcf2diem** function, which can be accessed directly from the command line and takes the vcf file name as an argument. The **vcf2diem** function performs a lossless compression of the data, simultaneously formatting and filtering it for use in `diempy`. Not all vcf file variants are passed to *diem*. To be accepted, the two most common alleles (A,B) of a variant must both appear in the homozygous state in at least two individuals in the dataset. Within accepted variants, genotype encodings involving rarer alleles give rise to ‘U’ genotypes within `diempy`, that is: unencodable with respect to (A,B) but still permissible for *diem*. For each accepted variant, the number of alleles, and whether those alleles occupy one base or a variable number of bases (indels) is recorded in the *diem input*.*bed* file. Rejected variants are recorded in the *diem excluded*.*bed* file with the reason for their exclusion. The **vcf2diem** data compression is lossless up to the ten most common alleles of a variant.

The third action of **vcf2diem** is to output a *diem meta* file. Each row corresponds to a chromosome in the dataset. Columns record the start and end base pair reference positions of chromosomes (thus, for most references, their length), the location of the first and last variants used for polarization, the total number of variants retained per chromosome, and their relative recombination rates (defaults to 1.0). Each remaining column specifies an individual and its *ploidy* with respect to each chromosome. In combination with the generalised homogeneous/heterogeneous *diem* state definitions [3] this allows arbitrary ploidies to be encoded (e.g. multiple hemizygous sex chromosomes or a mix of sexes sampled in a haplodiploid system). Because vcf files do not contain such meta information, by default, **vcf2diem** sets all ploidies to 2. While the *diem* algorithm is relatively robust to errors in ploidy, corrected ploidies should be entered by the user. For example, in an X-Y system, males are haploid for the X chromosome, so their ploidy for the X should be set to 1. To correct ploidy information, the user must supply a tsv file in which the first column lists individuals in the dataset (header *#Inds*) and each remaining column corresponds to a chromosome which has non-default ploidy (headers must match chromosome names in the original vcf file). From within `diempy`, the meta data file can be updated using the `diem.update meta()` function (the online tutorial provides an example).

### Polarization

To perform polarization, the data are loaded into `diempy` as an instance of the `DiemType` class (see implementation section above). Internally, polarization occurs in two steps. First, the data are flipped to a random null polarity. This removes any biases about the assignment of alleles to a parent population or species. For example, genotype calls in a vcf file are generally polarised with respect to the allelic state of the reference genome which may be an arbitrary individual haplotype or a consensus sequence across many individuals, a distant reference with no detailed statewise relevance to the analysis, or a reference constructed from one of the genomes being analysed. An example of a null polarity can be seen in Fig. 2A. Under null polarisation, the estimated hybrid index of each individual will be approximately 0.5. Second, the *diem* algorithm seeks to maximize the difference between individuals by placing them on either side of a *barrier* to gene flow identified as the biggest step in estimated hybrid index of the (sorted) individuals. It does so by iteratively checking the labeling of each variant, flipping the label when it increases the likelihood of the data given the current barrier position. The algorithm is deterministic in that a given null polarity (which can be input) results in one and only one polarisation.

**Figure 2:**
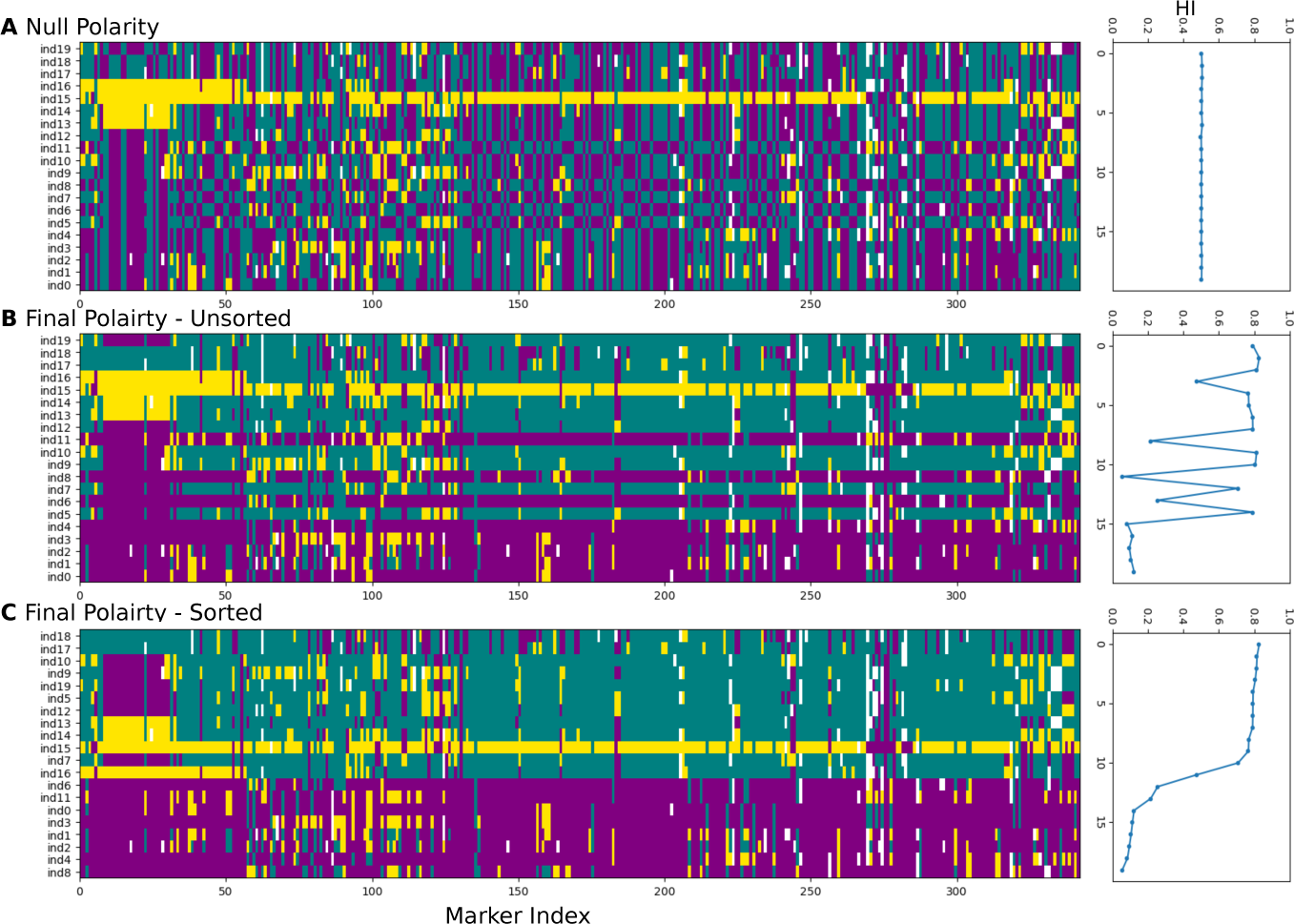
Polarizing variant data. The state of each individual is plotted for each variant indexed along the genome (i.e. the physical distance between variants is ignored). Purple and cyan correspond to the homozygous allele assignment that distinguishes the two sides of the barrier. Yellow corresponds to heterozygous states, and white corresponds to the U state (missing, or involving rare variant). The hybrid index of each individual is plotted to the right of the painting. A) the null polarity with alleles randomly assigned. B) the genotype states after polarizing. C) the polarised data with individuals sorted by *diem*-estimated hybrid index.

The output is a maximum likelihood labeling of the data that we visualize in Fig. 2B. We can distinguish the barrier more clearly by re-sorting the individuals by hybrid index (Fig. 2C). The polarised data can be stored directly as a `DiemType` object, which is efficient and useful for further analyses within `diempy`, or as a bed file which requires more memory but allows access for downstream analysis outside of `diempy`.

### Thresholding

For polarised data, `diempy` returns the Diagnostic Index (DI) for each variant, a measure of how well its configuration matches the identified genetic barrier. A variant configuration is simply a vector of genotypic states across individuals, corresponding to a given column of the painting in Fig. 2C. Note that many variants in the dataset will have the same configuration. We first consider the DI distribution, placing the set of unique configurations in the example dataset into 500 bins (Fig. 3A, cyan). The right-most bin contains the unique variant configuration that best fits the barrier and has highest DI, i.e. all individuals to the left of the barrier are homozygous for one allele while all individuals to the right of the barrier are homozygous for the alternative allele. In magenta, we show the counts of the variants in each of the cyan bins. For example, the most diagnostic variant configuration appears 10 times in the dataset.

**Figure 3:**
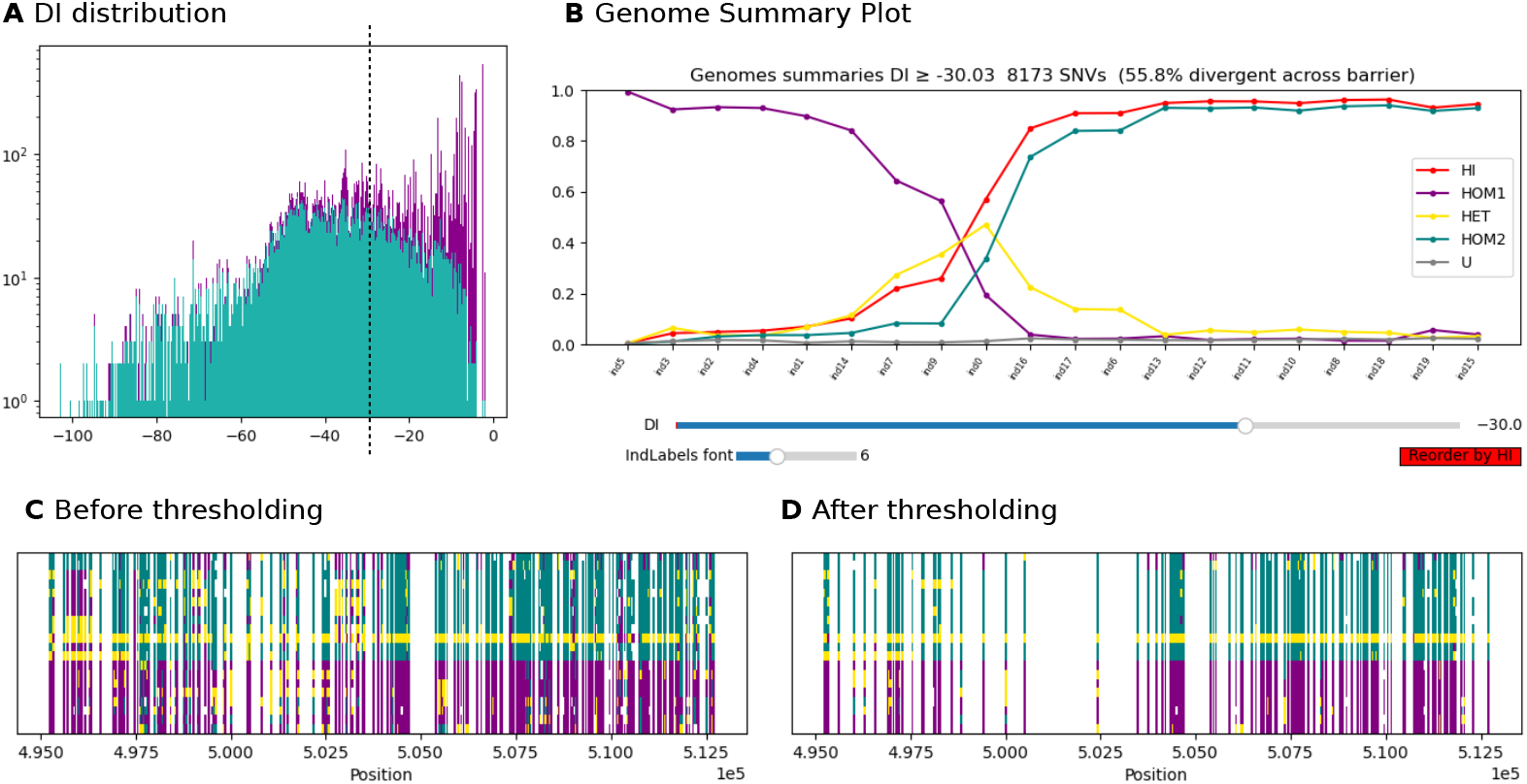
Applying a DI Threshold. A) the distribution of Diagnostic Index (DI) values. Cyan shows the counts of the unique variant configurations in 500 bins of DI values, with high values (closer to 0) corresponding to high diagnosticity. For the set of variant configurations in a given bin, the magenta bar indicates how many times those configurations appear in our dataset. B) four summaries of the data for each individual: the HI value, the proportion of ‘U’ (Unencodable) genotypes, and the proportion of genotypes homozygous for the two alternative alleles. C) the same data from Fig. 2 with variants in their physical positions and separated by whitespace. D) the variants remaining after removing sites with DI value ≤ −30.

Variant configurations with low DI values carry little information about the barrier that separates our samples and may correspond to ancestral shared variation or genotyping errors. Thus, it may be useful to remove those variants from downstream analysis. This can be achieved using the *threshold* function in `diempy`. The choice of threshold will be unique to each dataset and should be informed by the DI distribution and the research question. We provide the interactive `GenomeSummaryPlot()` (Fig. 3B) to explore how thresholding affects the data. For example, here we see that retaining variants with DI≥ −30 yields a strong separation, with the peripheral individuals achieving HI values close to 0 and 1. Of course, this threshold makes the data more sparse, as is evident in Figs. 3C and D, where variants are plotted at their relative position in the chromosome. It is worth remembering that even without DI thresholding, variant densities in population genomic data may be sparse and uneven.

### Smoothing

After thresholding, the polarised genotypes of an individual may consist of long blocks of variants with (mostly) identical state. In particular, we may be interested in tracts of admixture which, when rare, appear as consecutive runs of variants that are heterozygous by source (e.g. the central individual in Fig. 3C). The distribution of the lengths of these blocks is open to inference [5] and leads to very compact genome representations. However, on closer inspection, obvious blocks often contain tiny imperfections that would break up automated block length measurement. One solution is a low-frequency-pass filter that removes any high-frequency changes in state along the chromosome. We implement this using kernel smoothing.

We smooth the genotype of each individual separately. For a given individual, we trace along the chromosome and smooth each site independently. To smooth a site, we assign a weight to it as well as to the sites in its vicinity. We calculate the total weight of each possible state in this region, and the focal site is assigned the state with the highest weight. The state may change, or it may remain the same. Here, we use the Laplace Kernel to assign weights (Fig. 4A).

**Figure 4:**
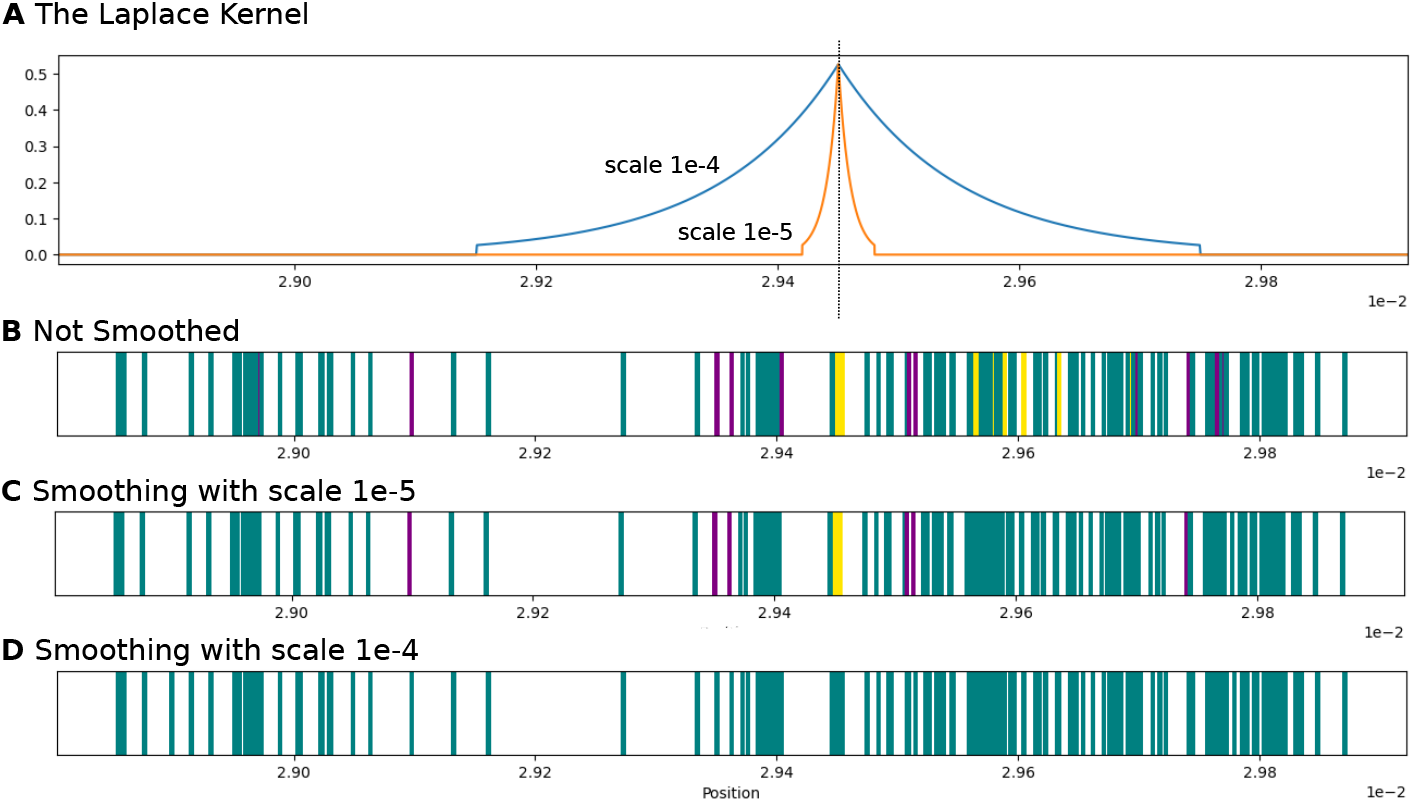
Smoothing polarised data with a Laplace kernel. The variants are plotted on a unit scale obtained by normalizing the base pair positions by the chromosome length. A) the (truncated) Laplace kernel centered on the indicated variant for two different scale values. B) the genotype of the second individual from the top in Fig. 3D. Panels C and D show the effect of smoothing this genotype using the two different scales.

The Laplace Kernel is a reflected exponential distribution and a natural choice: recombination diminishes the genealogical association between sites in the genome, and the strength of association between two sites breaks down exponentially with distance. It is therefore useful to think of the positions of sites on a unified scale that reflects the recombination rate. To do this, we normalize the base pair position of the sites by the total length of their respective chromosome, placing them along a unit interval. The motivation is this: if we assume that every chromosome must undergo a single recombination event in meiosis, the total recombination rate on each chromosome should be approximately the same. In the tutorial, we demonstrate how to modify this default to reflect organism-specific recombination dynamics (e.g. a difference in the scaled recombination rates between autosomes and sex chromosomes).

What remains is to choose the appropriate smoothing scale that determines the total breadth of the Laplace distribution. In Fig. 4A, we show the Laplace kernel with two different scales used to smooth the haplotype in Fig. 4B. For the smaller scale (1e-5), many of the small tracts have been smoothed over, but larger tracts, such as the one surrounding our focal heterozygous genotype, remain (Fig. 4C). By increasing the scale of the Laplace distribution, we start to smooth over larger tracts as well (Fig. 4D). Note that for computational efficiency, we truncate the Laplace distribution at a distance where the contribution of a given site becomes negligible.

In general, the appropriate choice of the Laplace scale will be unique to each dataset and research question. In the online tutorial, we provide guidance for testing and choosing among different scale values.

## Discussion

Above, we have illustrated the core function of `diempy`, i.e. genome polarisation across a genomic barrier. Population genomic data are informative both about long term barriers and recent admixture. `diempy` provides an entry point for exploring these data as well as additional tools for population genomic analysis. We briefly outline some of these below. We refer the users to the online tutorial for an in-depth tour.

### Analysing tract lengths

Once we have applied any DI threshold and/or smoothing to the polarised data, individual genomes can be represented as a set of tracts of contiguous ancestry. The length distribution of these tracts across a sample is highly informative, for example, about the rate and timing of admixture [2, 5]. To access this information directly, we apply the `create_contig_matrix()` function to our `DiemType` instance. This returns and a run-length encoding of genotypes as a contig – an ordered list of contiguous intervals of identical state. In the tutorial, we demonstrate how to construct contigs, plot the length distribution of ancestry tracts, and save this information in BED format.

### Browsing the Data

Genome polarisation data are typically very rich. We provide interactive visualisations that summarise data along multiple axes, making previously published styles of plot available publication-ready. Interactive ‘DI sliders’ allow the user to examine the consequences of different thresholding levels. A global view of all sites in the data forms a circular ‘iris plot’ which may show that barrier sites (maximum DI) are over-represented on certain chromosomes (e.g. sex chromosomes) or restricted to particular genomic regions (e.g. a polymorphic inversion in an otherwise unstructured population). Plots of polarised chromosomes may also reveal segregating introgression tracts and their shared boundaries, Fisher junctions [6, 7] across individuals. All feature plots are browsable: placing the cursor over a feature reports its genome location in reference coordinates (and the individual(s) in which it occurs).

### Masking

It is rare that a first analysis is final. An initial genome polarisation may reveal outlier genomes or genomic regions that are not relevant to the barrier of interest. Rather than restarting the *diem* workflow from scratch, **Masking** allows the user to refine their analyses mid-flow. Crucially: masked data are still polarised by *diem*, but have no influence over that polarisation. For example, an outlier individual incorrectly included from a sister taxon can be masked to remove its influence (without constructing a new vcf file), while being polarised to show potential introgression relevant to the barrier of interest.

On the along-genome axis, a region of low effective recombination (little change in polarised state across many sites) may correspond to a trans-species polymorphic inversion. While the inversion is its own (old) local barrier, the user may not wish to confound its effects with recent (speciation) barrier signal. Once the inversion is identified by an initial polarisation, it can be masked for an iterative *diem* analyses, which may reveal barriers associated with the more recent species divergence. Similarly, mitochondria and non-recombining Y,W chromosomes should be masked: polarisation will then highlight sites which differentiate variation in these non-recombining compartments across the focal barrier. Each system is likely to have its own optimal masking, but the path to do so is straightforward in the bed file workflow, and detailed in the tutorial.

### Future directions

Most analyses of population genomic variation are necessarily explorative. Even when we wish to test strong prior hypotheses, understanding genome scale variation is an iterative process that involves both discovery and the testing of hypotheses. We envisage `diempy` analysis as an iterative process of fast polarisation and browsable visualisation allowing the user to focus analysis clearly on their barriers of interest. We hope that the modular design of `diempy` will facilitate further development, both for the polarisation step and post-polarisation analyses. Firstly, the polarisation procedure can be generalised to simultaneously identify multiple barriers to gene flow among the samples, reducing the potential bias introduced by exploratory and iterative analyses. While it might be thought that focusing on two allelic states limits the number of barriers to two, it is the conformation space of allelic states across individuals that determines how many barriers can be distinguished. Secondly, it is straightforward to change the underlying smoothing kernel or implement alternative approaches to smoothing such as an HMM. Thirdly, an obvious downstream analysis post-polarisation is the inference of geographic or genomic clines. Finally, there is a clear connection between polarisation and the Ancestral Recombination Graph (ARG) [11] that represents the history of a sample. Each unique variant configuration of polarised data corresponds to a particular branch in the ARG that bipartitions the sample. Thus, in principle, one could infer an ARG and polarise this directly. It will be useful to quantify – for a known admixture history and ARG – to what extent particular DI values correspond to recent introgression vs shared ancestral variation.

## Data Availability

The source code for `diempy` is available on GitHub at https://github.com/Studenecivb/diemPy and is available for installation via PyPi at https://pypi.org/project/diempy/. The main documentation is provided as a comprehensive tutorial available at https://github.com/DerekSetter/tutorial-DiemPy/ with extensive API documentation (https://diempy.readthedocs.io/).

## Acknowledgments

We would like to thank Zia Leigh and Meng Lu for testing early versions of the software.

## Supplement

### Simulation

Simulations for benchmarking were performed using msprime [15, 4] v1.3.4 under a scenario of secondary contact between two highly diverged populations/species. We simulated 100 replicate datasets each consisting of a sample size of *n* = 1, 000 diploid individuals under a history of a single pulse of admixture 10 generations ago. Mutations are added under the infinite sites model, and the replicates are treated as individual chromosomes that are combined to create a single, large dataset with sufficiently many mutations to perform the benchmarking. In subsetting, we always select an equal number of individuals from each population and an equal number of mutations on each chromosome. The simulation code for a single iteration (chromosome) is as follows:

~~~
pSize = 1_000
splitTime = 10*(pSize*4)
timeSinceContact = 10 #generations
migRate = 1/pSize * 100
contactDuration = 1 #generations
muRate = 1.2e-6
recRate = 1e-7
seqLength = 1e6
nDiploidsSampled = 500 # 500 each population
demography = msprime.Demography()
demography.add_population(name = ‘anc’, initial_size= pSize)
demography.add_population(name = ‘A’, initial_size=pSize)
demography.add_population(name = ‘B’, initial_size = pSize)
demography.set_symmetric_migration_rate(populations=[‘A’,’B’],rate = 0)
demography.add_symmetric_migration_rate_change(
    time = timeSinceContact,rate = migRate, populations=[‘A’,’B’])
demography.add_symmetric_migration_rate_change(
    time = timeSinceContact+contactDuration, rate = 0, populations=[‘A’,’B’])
demography.add_population_split(
    time = splitTime, derived = [‘A’,’B’], ancestral = ‘anc’)
ts = msprime.sim_ancestry(
    samples = *{*‘A’:nDiploidsSampled,‘B’:nDiploidsSampled*}*,
    demography=demography,
    recombination_rate=recRate,
    sequence_length=seqLength,
    discrete_genome= True
)
mts = msprime.sim_mutations(ts,rate=muRate,discrete_genome=False)
~~~

